# Probing the neurophysiology of temporal sensitivity in the somatosensory system using the mismatch negativity (MMN) sensory memory paradigm

**DOI:** 10.1101/2023.06.21.545720

**Authors:** Emily L. Isenstein, Edward G. Freedman, Ashley J. Xu, Ian A. DeAndrea-Lazarus, John J. Foxe

## Abstract

Duration is an amodal feature common to all sensory experiences, but current understanding of sensory-perceptual processing of the temporal qualities of somatosensation remains incomplete. The goal here was to better understand how the brain processes the duration of vibrotactile information, which was assessed by parametrically varying the extent of duration deviance in a somatosensory mismatch negativity (sMMN) paradigm while high-density event-related potential (ERP) recordings were acquired. Healthy young adults (N = 20; aged 18-31 years) received stimulation of the right index fingertip with a 100 ms vibro-tactile input on 80% of trials while the other 20% of trials consisted of deviant stimuli with one of the following durations: 115, 130, 145, or 160 ms. Deviant conditions were presented in separate blocks with deviants pseudo-randomly distributed amongst the 100 ms standards. Participants ignored these inputs while watching a silent movie. Robust sMMN responses, with a dipolar field over the left antero-superior parietal cortex, were detected when deviant stimuli were 130, 145, and 160 ms, but not when they were 115 ms. The amplitudes of the sMMN correlated with individuals’ subsequent abilities to detect duration deviants when actively attempting to discriminate their presence. This simple-to-execute sMMN paradigm holds promise for the assessment of tactile processing differences in clinical populations where tactile sensitivities are a common aspect of the phenotype (e.g., Autism, Fragile-X syndrome).

## INTRODUCTION

Temporal precision of the somatosensory system is critical for the discrimination of tactile stimuli, heavily influencing the perception of material properties. The integration of physical features over time provides insight into details such as texture^1^, orientation^2^, and spatiality^3^, allowing for the formation of dimensional representations of objects. While the threshold of temporal discrimination varies depending on stimulus parameters such as posture and movement^4,5^, research has shown that humans can distinguish the temporal order of sensory stimuli with a fidelity of approximately 30 ms^6,7,8,9^. While somatosensory temporal precision has been extensively explored behaviorally, much less is known about the electrophysiological signatures of this processing capacity. Exploration of the relationship between the detection of a temporal discrepancy with the degree of electrophysiological response carries potential to elucidate what drives atypical sensory sensitivities in certain clinical populations.

One very well-characterized probe of temporal processing, albeit mainly in the auditory system, is the event-related potential (ERP) component known as the mismatch negativity (MMN), which captures the brain’s ability to detect deviations from a sensory expectation^10^. By maintaining one input in a sequence as the most probable, any perceived deviation from the anticipated stimulation results in a distinct negative electrophysiological deflection known as the MMN. The MMN is also pre-attentive, meaning that deviance is detected automatically and active engagement with the stimuli (i.e., directed attention) is not required to elicit the electrophysiological signature^11-15^. As such, the ability to collect this measure passively allows for the assessment of populations who may not be able to complete active tasks, such as those with intellectual disability^16,17^, Fragile X Syndrome^18^, schizophrenia spectrum disorder^19-21^, Rett Syndrome^22^, and autism spectrum disorder (ASD)^23-26^.

While mismatch negativity is most often elicited by auditory (aMMN)^10,27^ or visual stimuli (vMMN)^28,29^ due to ease of presentation, the MMN can be produced across many different contexts. The MMN can be produced by varying many feature dimensions of a stimulus stream, such as frequency^30-32^, physical location^33-36^, and duration^32,37-39^ of the stimulus, demonstrating the robustness of the signal. The somatosensory MMN (sMMN) has also been established as a reliable measure of deviance detection and has been elicited with multiple different modes of tactile stimulation (e.g. vibrotactile^33^, electrical^34^, compressed air^36^), to study tactile processing in infants^36^, children ^33^, as well as young and elderly adults^40^.

For example, the sMMN has been used to investigate the sensitivity of the somatosensory system to differences in the duration of vibrotactile stimuli in multiple experimental paradigms comparing standard and deviant vibrations^32,37-39^. Intracranial recordings have been used to localize the activity to somatosensory regions of the post-central gyrus, further validating the effectiveness of this paradigm to assess low-level tactile processing^37,41^. These studies examined temporal sensitivity in the somatosensory domain across a broad range of durations, yet a systematic comparison of the electrophysiological responses to parametrically varied differences in tactile duration has not been conducted. As such, the sensitivity of this system to duration differences has not been fully detailed.

The aim of the present study was to investigate how the degree of deviance in the duration of somatosensory input would affect the sMMN signal in neurotypical young adults. With greater understanding of somatosensory perception, this paradigm can be further applied to study the cortical underpinnings of atypical tactile sensitivities in other groups. For example, hyper-sensitivity to tactile input is common in ASD and can cause significant pain and distress^42^, yet the neurocognitive underpinnings of this sensitivity remain unclear. To achieve this aim, an auditory MMN paradigm was adapted to evaluate duration discrimination along the same parameters in the somatosensory domain using high-density electroencephalography (EEG)^43^. This methodology allowed us to investigate cortical deviance detection to a series of vibrotactile duration changes, probing low-level somatosensory discrimination.

## METHODS

### Participants

The study included 20 adults aged 18 to 31 (mean 24.03, STD 3.59; 13M/7F). All self-reported normal hearing and normal or corrected-to-normal vision. No participant reported a history of traumatic brain injury, neurologic disorder, autism spectrum disorder, schizophrenia, or psychosis. All gave written informed consent and were paid an hourly rate for their participation. All procedures were reviewed and approved by the Research Subjects Review Board (RSRB) at the University of Rochester (STUDY00002036).

### Stimuli and Task

Participants sat in an electrically-shielded and sound-attenuating EEG booth (Controlled Acoustical Environments, North Aurora, IL, USA). Somatosensory stimulation was administered via an Adafruit 1201 vibrating mini-motor disc (Adafruit Industries LLC, New York, USA) connected to an Arduino Uno microcontroller (Arduino, Turin, Italy). The mini-motor disc was secured around the participant’s right index fingertip with Velcro such that the disc was snugly secured, but not so tightly that the participant began to feel their own pulse. Stimulus presentation was controlled by Presentation® software (Version 18.0, Neurobehavioral Systems Inc., Berkeley, CA, USA) and all vibrations were administered at a voltage of 5 V and a frequency of 183.33 Hz. Participants wore either noise reducing ear muffs (n = 5; MPOW HM035A; Longgang, Guangdong, China) or foam earplugs (n=15; Laser Lite LL-1, Howard Leight by Honeywell, Charlotte, North Carolina, USA) to rule out air-conducted hearing of the vibrating mini-motor disc, and they were instructed to ignore the vibrotactile stimuli during the electrophysiological recording component of the experiment. During EEG data collection, they watched a silent video of their own choosing with optional subtitles. There was no video presented during the subsequent behavioral component of the task. Participants rested their right arm comfortably on a table in front of them with their palm oriented upwards. They were requested not to move their right hand and not to rest their right index finger on the table throughout the experiment. It is important to note that participants were not informed of the purpose of the experiment before or during the initial electrophysiological stage of the experiment; that is, they were not informed that there were “duration deviants” in the blocks of presented stimuli. This was only revealed to them before the behavioral portion of the experiment, which always followed the EEG portion.

There were four condition blocks presented in a random order to each participant in which 80% of stimuli presented were the standard duration and 20% were the deviant duration. Each block presented the same standard stimulus (100 ms) but used a different duration deviant stimulus (115, 130, 145, or 160 ms, respectively). A total of 1000 stimuli were presented per block in a pseudo-random order per block, with no two deviants presented consecutively. Stimuli were presented with an average inter-stimulus interval of 750 ms with a random temporal jitter between ±150 ms.

### Electroencephalographic (EEG) Recordings

EEG was recorded using a 128-channel BioSemi high-density EEG system (Biosemi ActiveTwo, Amsterdam, The Netherlands). Continuous data were recorded at sampling rate of 512 Hz and low-pass filtered during collection at 1/5 this rate, as per the default Biosemi decimation filter, then processed with MATLAB 2021a software (Mathworks, Natick, MA, USA) and the EEGLAB toolbox (version 14.1.2)^44^ using custom scripts. Data were deemed of acceptable quality in the unfiltered state; as such, the remaining pre-processing steps were conducted without the application of any further digital filtering. Channels with amplitudes ± 3 standard deviations from the mean were removed and interpolated. The data were then re-referenced to the average reference and data were aligned on stimulus presentation and epoched from -100 to 500 ms. Epochs were baseline corrected to the 100 ms prior to stimulus onset. Automatic artifact rejection criteria were used based on the normal distribution of the maximum and minimum values. Trials with amplitudes > 200 μ V were removed. As consistent with the EEG electrode locations identified in previous literature^30,32,38^, the MMN was extracted from the average of a cluster of fronto-central electrodes (illustrated in Figure 3). Topographic maps of the voltage distributions across a 2-dimensional circular representation of the scalp were produced using interpolation projected on a cartesian grid from the “topoplot” function in the EEGLAB toolbox in MATLAB.

### Behavioral Discrimination Task

After completing the EEG portion of the experiment, participants completed a shorter behavioral version of the task that served as a discrimination task to assess perceptual thresholds. This behavioral task had the same stimuli ratio but included only 100 total stimuli per block. Stimuli were divided into four blocks (consisting of the four different duration deviants) and the order of presentation was randomized across participants. Each block contained 80 standard and 20 deviant stimuli. Participants were instructed to attend to the stimuli and press a button with their left hand whenever they detected a deviant stimulus.

### EEG Data Analyses

The point of maximum difference between the deviant and respective standard response was calculated between 150 and 250 ms post-stimulus to assess the latency of the MMN for each individual participant, and this was individually determined for each duration-deviant condition. These individual latency values were then averaged to determine the points of maximum difference on a group level for each duration condition, leading to the following values: 115-dur = 185 ms; 130-dur = 178 ms; 145-dur = 183 ms; 160-dur = 194 ms. The average MMN latency across all four conditions was 185 ms. A 50 ms window (160-210 ms) was defined around this peak for all subsequent calculations of MMN amplitude, which was defined as the integrated amplitude across this 50 ms time-bin. A two-way repeated measures analysis of variance (rmANOVA) was conducted to test for effects of condition (standard vs. deviant) and duration deviance (115, 130, 145, 160 ms) using these amplitude measures, consistent with prior work^43^. A one-way repeated measures ANOVA was used to test for latency differences across duration deviance conditions. Difference waves were calculated by subtracting the individual standard waveforms from each respective deviant waveform for visualization purposes.

We calculated d-prime (d’) measures of sensitivity to deviance detection and conducted a one-way repeated measures ANOVA to analyze the effect of duration deviance (115, 130, 145, 160 ms) on the d’ value. Pearson correlation was also calculated between each behavioral d’ in response to the four deviant durations and the respective MMN amplitudes; each participant contributed four points to the correlation, one for each duration deviance.

### Post-Hoc Exploratory Analyses

In order to more fully explore the rich spatio-temporal dynamics of this high-density ERP data matrix, a secondary exploratory analysis phase was also conducted. Statistical cluster plots were produced to visualize the localization and strength of differences in scalp topography between standard and deviant stimuli. Plots were generated from a cluster-based permutation test using a series of two-tailed, independent sample t-test (α = 0.05) using the FieldTrip toolbox in Matlab. Monte-Carlo sampling was applied using 5000 iterations to model outcome probability, triangulate neighboring nodes for spatial clustering, and correct for multiple comparisons.

## RESULTS

### Behavioral Data

The behavioral detection sensitivity d’ values for each of the duration-deviance conditions were as follows: for the 115 ms d’ = 0.38 (Standard Deviation (SD) = 0.49; Range (R) = -0.96 - 1.20); 130 ms d’ = 0.80 (0.72; -0.46 - 2.24); 145 ms d’ = 0.97 (SD = 0.95; R= -0.55 - 2.92); 160 ms d’ = 2.06 (SD = 1.03; R = 0.36 - 4.48). Within the more difficult, shorter duration deviant conditions, there were a number of participants who did not correctly detect any deviant stimuli, with 5 (of 20) in the 115-deviant condition, 2 in the 130-deviant condition, and 5 in the 145-deviant condition. All participants were able to detect deviants in the 160-deviant condition (see **Figure 1, top panel, hit rates)**. The rmANOVA yielded a significant main effect of the degree of deviance (F(3, 57) = 25.81, p = 1.12^-10^, η^2^ = 0.58). Sphericity was not violated (p = 0.60). Post-hoc paired comparisons revealed that the 115 ms d’ was significantly smaller than the 130 ms d’ (p = 0.01, 95% C.I. = -0.87, -0.11), 145 ms d’ (p = 4.3^-3^, 95% C.I. = -1.06, -0.23) and the 160 ms d’ (p < 8.08^-6^, 95% C.I. = -2.13, -1.28), while the 130 ms d’ was significantly smaller than the 160 ms d’ (p = 3.09^-5^, 95% C.I. = -1.69, - 0.73) and the 145 ms d’ was significantly smaller than the 160 ms d’ (p = 4.0^-6^, 95% C.I. = -1.40, -0.72). All other p’s were greater than 0.49.

**Figure 1.**
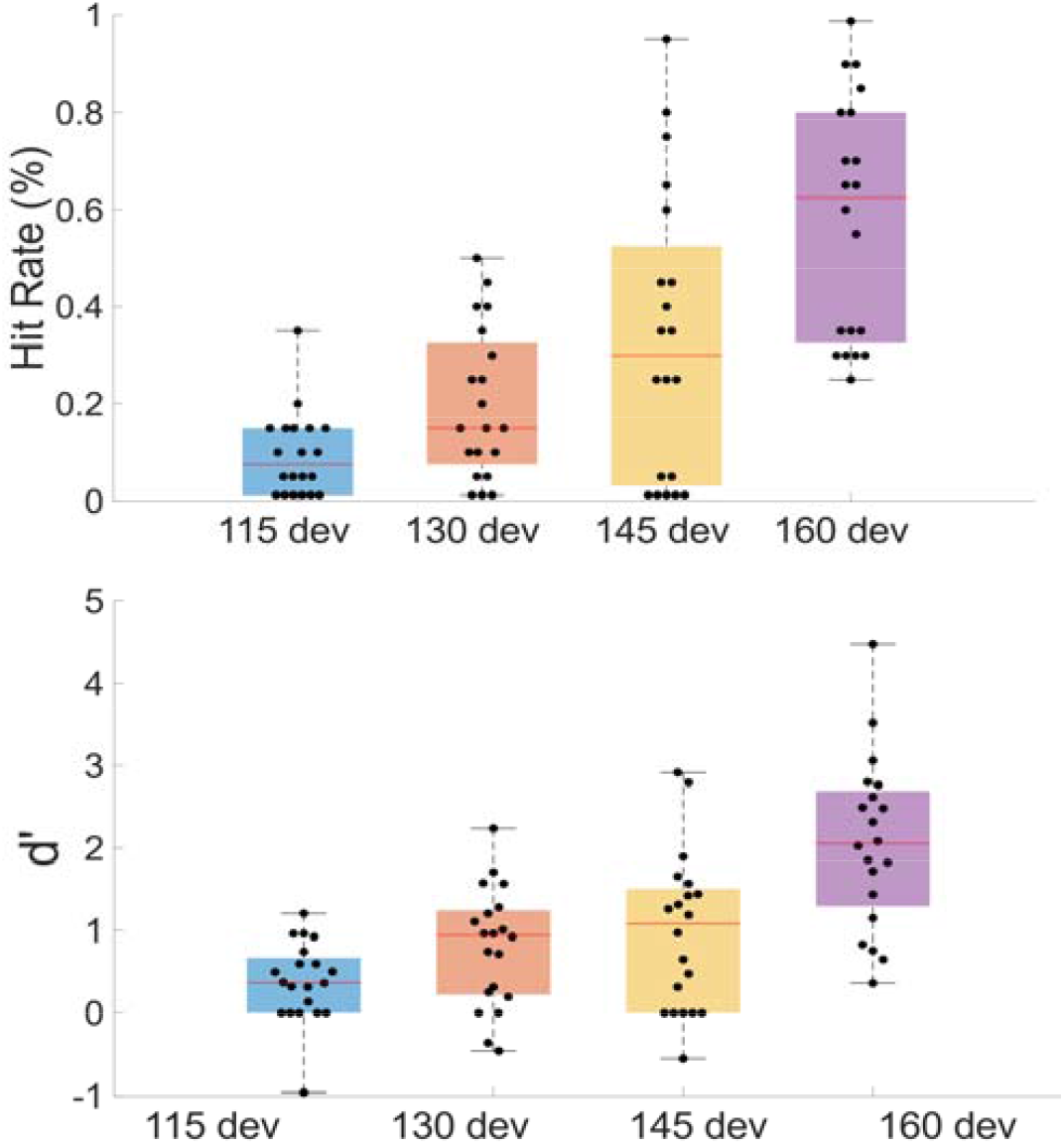
**A)** Percent hit rate for each duration condition. **B)** d-prime value for each duration condition.

### Electrophysiological Data

Grand averages of event related potential (ERP) waveforms for the standard and deviant conditions are plotted in **Figure 2A**. The difference waves for the deviant conditions and the distributions of the peak MMN amplitudes are plotted in **Figure 2B and Figure 2C**, respectively. Group means for amplitude and latency are presented in **Table 1**. Several earlier sMMN studies also found a positive deflection following the MMN response^32,34,37,39,41^; Visual inspection of the current data set suggested no evidence of this positive deflection following the MMN so group means for amplitude and latency were not calculated in this domain. Individual level data on the average amplitude between 160-210 or for the standard and deviant conditions can be found in **Figure 3**.

**Figure 2.**
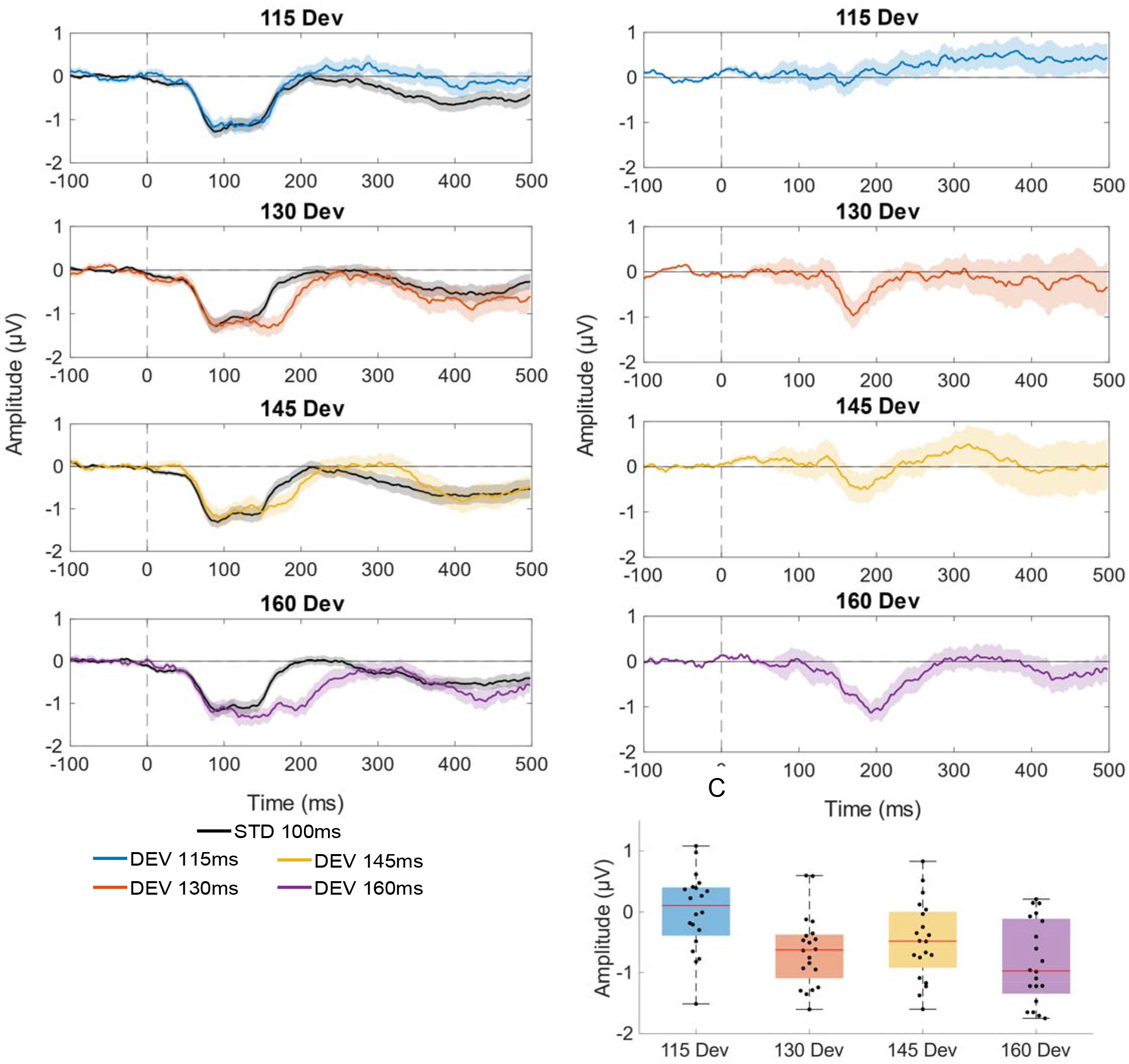
**A)** Grand average ERPs for the standard (100 ms) and deviant (115, 130, 145, 160 ms) vibrations. **B)** Distribution of peak MMN difference wave amplitudes for the 115, 130, 145 and 160 ms deviant conditions. **C)** Grand average MMN difference waves for the 115, 130, 145 and 160 ms deviants.

**Figure 3.**
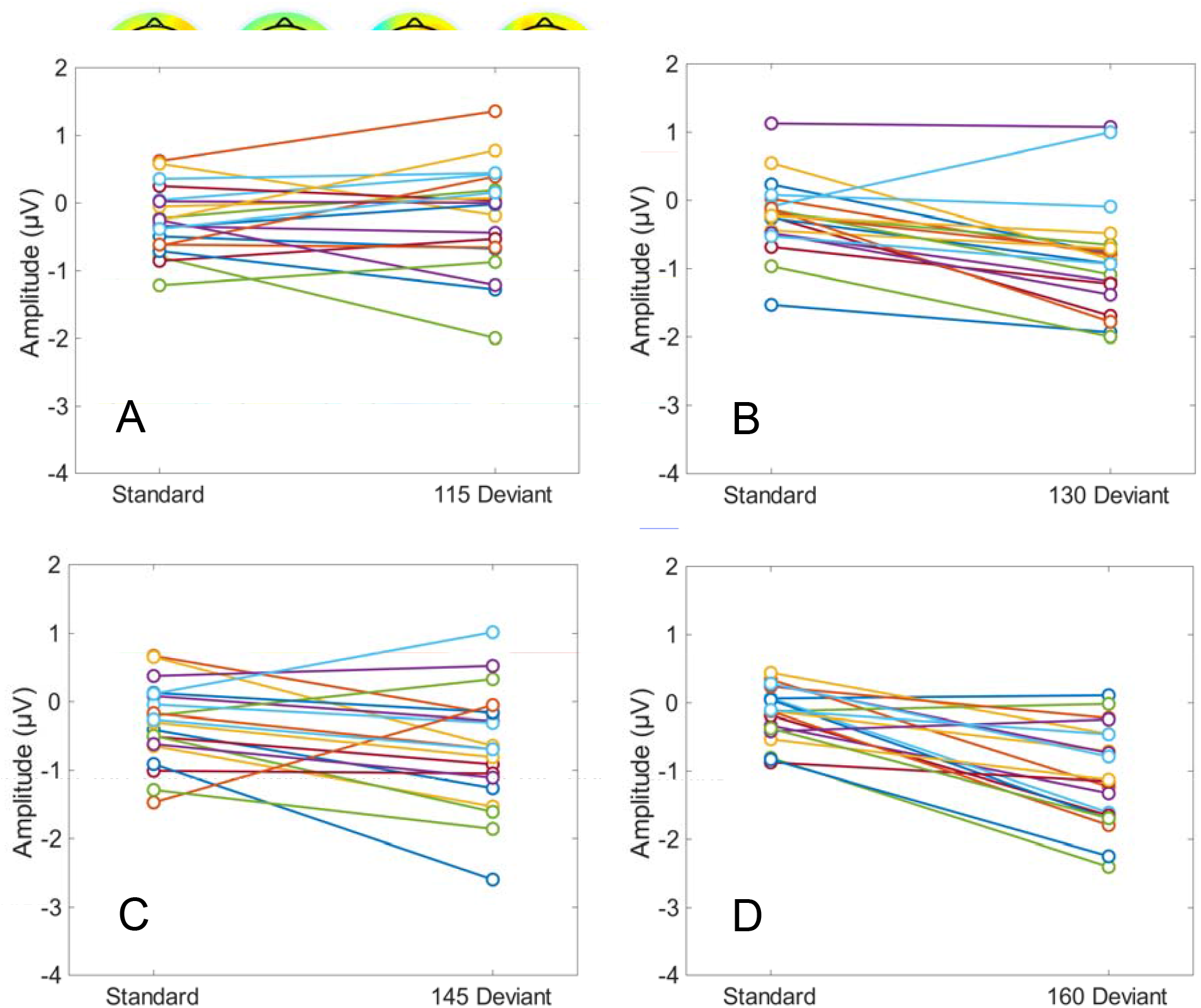
Line plots displaying individual participant change in amplitude between the standard and 115 (**A)**, 130 (**B**), 145 (**C**), and 160 (**D**) ms deviant conditions.

**Table 1.** Mean amplitudes and latencies of 115, 130, 145 and 160 ms deviant MMNs (SE).

|  | Amplitude Group Mean (SE) | Latency Group Mean (SE) |
| --- | --- | --- |
| 115 Deviant | 0.06 $\mu\text{V}$ (0.13) | 184.9 ms (7.78) |
| 130 Deviant | -0.63 $\mu\text{V}$ (0.13) | 178.2 ms (4.28) |
| 145 Deviant | -0.38 $\mu\text{V}$ (0.16) | 182.7 ms (4.36) |
| 160 Deviant | -0.92 $\mu\text{V}$ (0.14) | 194.1 ms (4.89) |

Two way rmANOVA revealed a significant interaction between the effects of condition and duration deviance (F(3,57) = 10.05, p = 2.10^−5^, η^2^ = 0.35), and a main effect of both condition (F(1,19) = 28.08, p = 4.10^−5^, η^2^ = 0.60) and duration deviance (F(3,57) = 4.47, p = 6.93, η^2^ = 0.19). Sphericity was not violated for any measure (p’s greater than 0.36). Post-hoc planned comparisons revealed that the value of mean 115 MMN amplitude was significantly different than the 130 ms (p = 0.02 95% C.I. = 0.05, 0.57), the 145 ms (p = 0.01, 95% C.I. = 0.07, 0.58) and the 160 ms deviant conditions (p = 3.57^-3^, 95% C.I. = 0.14, 0.62). All other p’s were greater than 0.38.

One-way repeated measures ANOVA revealed no main effect of condition on latency, with no statistically significant differences in MMN latency (F(3, 57) = 1.53, p = 0.22, η^-2^ 0.07). Sphericity was not violated (p = 0.21).

The topographies of the MMN difference waves (i.e. deviant minus standard) for each of the four deviant conditions are displayed over the critical time-period (100 – 220 ms) in the plots in **Figure 4**. Clear dipolar activity is visible in the 130, 145, and 160 ms deviant conditions, but not in the 115 ms deviant condition, mirroring the demonstrations of the MMN waveforms in **Figure 2**. The absence of any notable topographic activity in the 115 ms condition underscores the finding that the brain’s responses to the 100 ms standard and the 115 ms deviant vibrations are not detectably different during the MMN processing timeframe. This pattern is consistent with the results of the above ANOVA, where there was no evidence for a detectable MMN in the shortest duration condition (i.e. to the 115 ms deviant). The variability in the strength and longevity of the dipoles also reflects the characteristics of the ERPs, with weaker, but present, MMN activity in the 145 ms deviant condition, and extended duration in the 160 ms deviant condition.

Pearson correlation between each individual’s deviant MMN amplitudes and the corresponding d prime values was -0.41 (p = 1.93^-4^) and is plotted in **Figure 5**.

**Figure 5.**
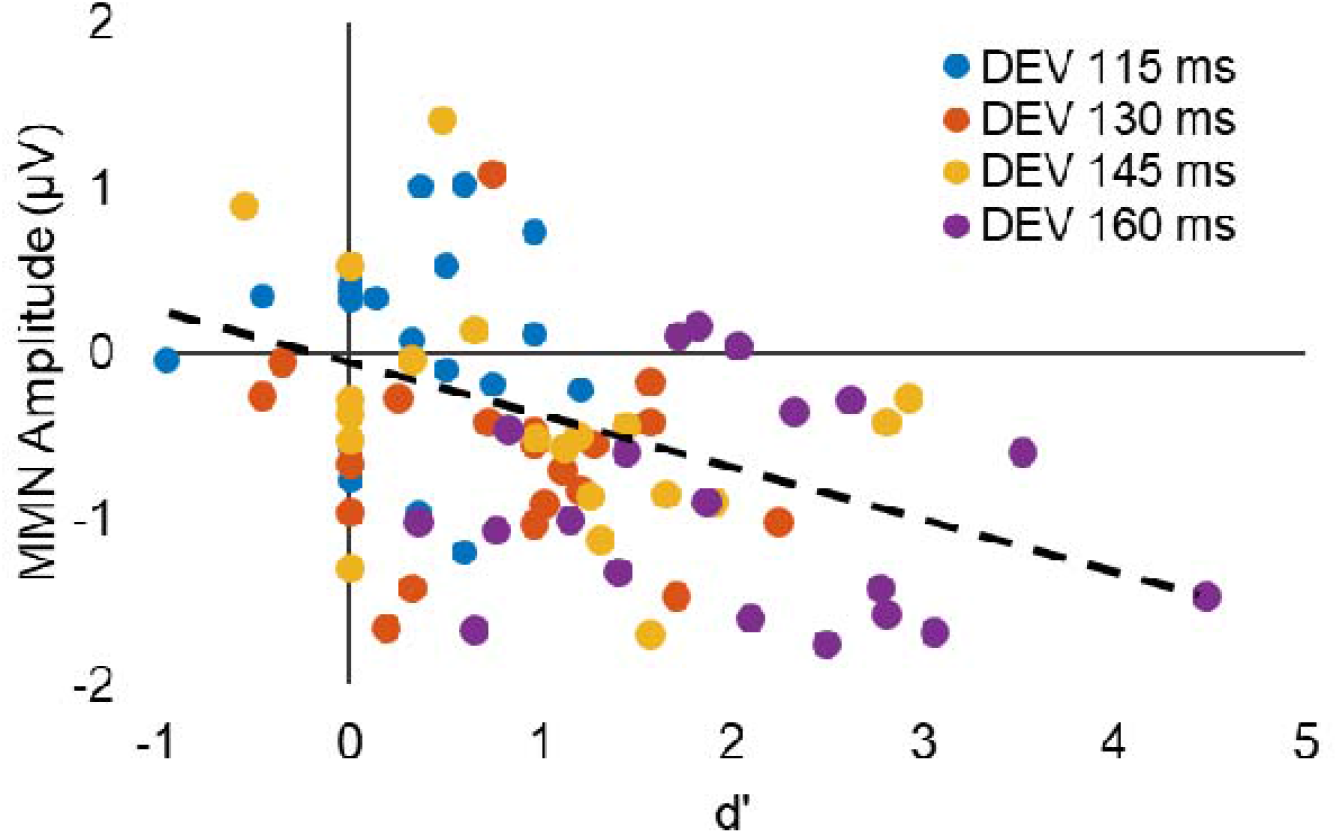
Pearson correlation between the behavioral d’ and MMN amplitudes for each the four deviant conditions.

Statistical cluster plots in **Figure 6** provide a comprehensive detailing of periods of significant difference between standards and deviants across the entire data matrix (i.e. all electrode sites X all time points (0 - 400 ms). Here again, evidence for significant differences during the MMN timeframe is clear in the three longer duration deviance conditions, but is absent in the shortest. The time-scale of the significant activity at approximately 150-250 ms post-stimulus recapitulates the focused time scale of the MMN, and clusters indicating a significant difference between the standard and deviant conditions are most prominent at the front-central and central-parietal areas.

**Figure 6.**
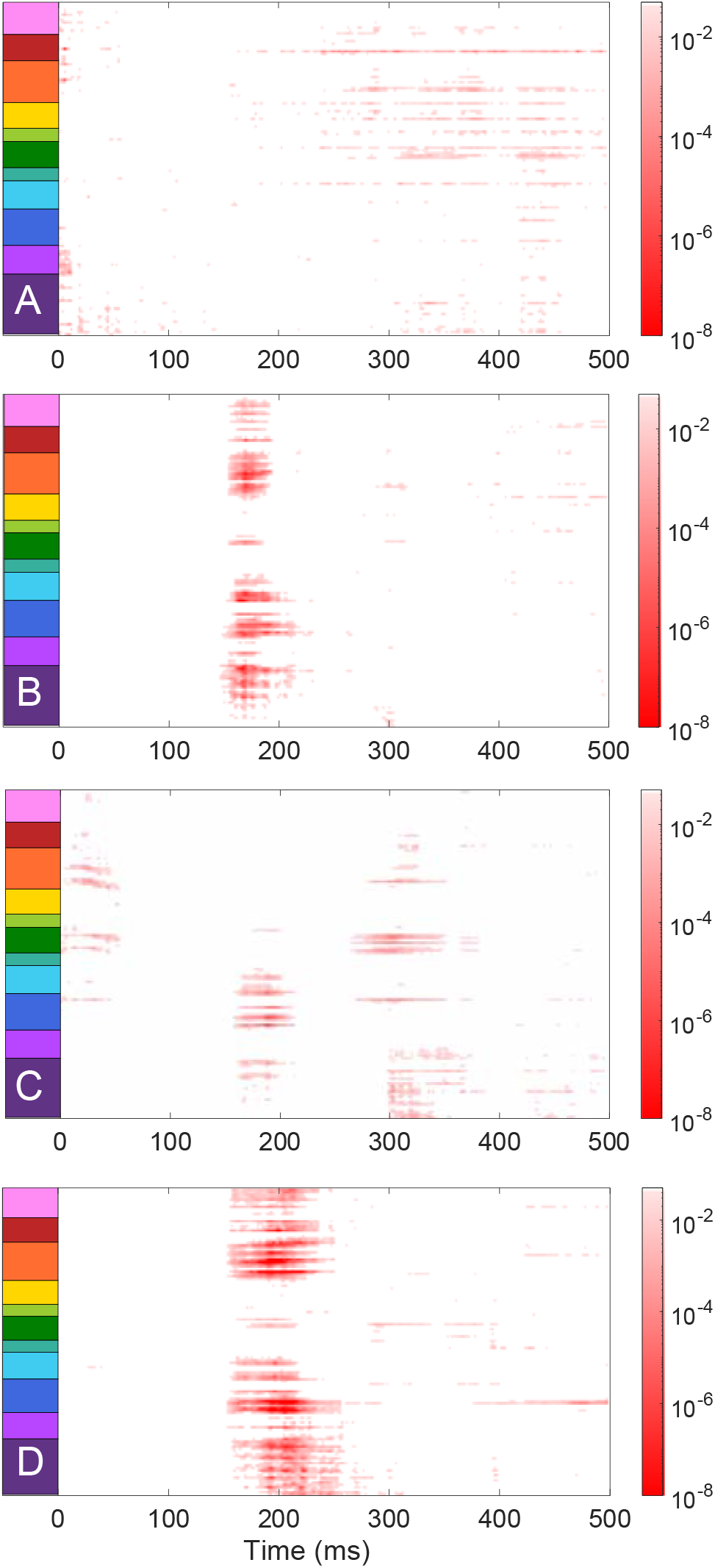
Statistical cluster plots representing significant ERP amplitude differences between standard (100 ms) and deviant (**A**. 115, **B**. 130, **C**. 145, and **D**. 160 ms) conditions. Two-tailed independent sample t-tests were used to calculate the p-values, with red indicating p < 0.05 and white indicating p > 0.05. The y-axis represents clusters of electrodes sorted by location and displayed here.

## DISCUSSION

The current investigation of duration somatosensory mismatch negativity yielded results consistent with previous studies that tested a similar somatosensory paradigm, with a negative deflection between 150 and 250 ms post-stimulus^32,38,41^. At the group level, statistically detectable MMNs were elicited when the deviants were greater than the 100 ms standard vibrations by 30, 45, and 60 ms, but not by 15 ms. Follow up analyses showed that the amplitudes of the MMN, measured while participants were ignoring the stimulus trains, were associated with individuals’ subsequent abilities to detect the deviants when they were asked to actively discriminate their presence in a follow-up behavioral paradigm. The substantial correlation between the behavioral d’ values and the MMN amplitudes points to an association between one’s ability to detect differences in the duration of vibrotactile stimuli both consciously and unconsciously, bolstering decades of work demonstrating a strong link between MMN activity and discriminative abilities^11-15^.

These findings align well with previous work that placed the threshold of vibrotactile discrimination at approximately 30 ms^6,7,8,9^, corroborating this paradigm as a measure of the neural correlates of temporal somatosensation. Further, the lack of differences in the latency of the MMNs indicates that the MMNs were not delayed in response to longer deviant vibrations. Accordingly, the differences in MMN amplitude are unlikely to be attributable to varying time scales of the responses to the deviant vibrations.

Unlike some prior studies^32,34,37,39,41^, the current paradigm did not elicit an obvious positive deflection following the negative MMN shift. One possible explanation for this difference is that prior studies utilized fixed inter-stimulus intervals, whereas the present study utilized a variable inter-stimulus interval. As such, the vibrations in the current experiment were less predictable and may have elicited a different electrophysiologic profile, suggesting that this positive component may reflect a fulfilled expectation that is not produced by the current paradigm.

The dipolar pattern seen in the topographic plots provides additional support for a parietal source of sMMN activity, which aligns with the intracranial results of Butler’s 2011 and Spackman’s 2010 papers that identify the somatosensory cortex as the source of sMMN activity induced by vibrotactile stimulation^37,41^. By targeting the cortical processes of low-level somatosensory processing, there is potential to investigate variability in response to tactile stimulation in clinical populations. Tactile hyper-sensitivity is common in conditions such as Fragile X Syndrome and autism spectrum disorder, yet the origin of this atypical sensory processing remains poorly characterized^42,45-47^. As mentioned above, a key feature of the MMN paradigm is that it does not require active engagement by the participant, and in this regard, it presents as an ideal way to non-invasively assay tactile sensitivity in vulnerable populations.

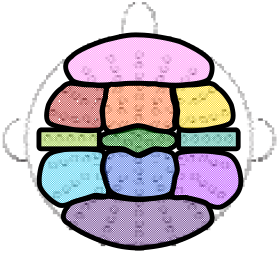

## Limitations

While the present study utilized duration as the deviant feature, the nature of the stimulus presentation method meant that vibrotactile intensity levels may have differed slightly between individuals. Because the MMN involves calculating the difference wave rather than the overall amplitude, the internal consistency of intensity (i.e. standards and deviants are of the same intensity) allows for effective acquisition of the magnitude of deviance detection at an individual level. The differing degrees of attention between the EEG and the behavioral tasks also does not allow for a direct comparison between the results. Numerous instances of a 0% hit rate in the 115, 130, and 145 ms deviant conditions of the behavioral task also indicate that even when the stimuli are attended to, the short differences in duration can be difficult to detect. A follow-up study that records electrophysiological activity while the participants attend to the vibrations would be beneficial to understanding the role of attention. Given the 15 ms increments in deviant duration, the fidelity of deviance detection between 15 and 30 ms in these conditions remains unknown and may also be investigated in future work. While the amplitude of the MMN was significantly different between the 115 ms deviant and each of the 130, 145, and 160 ms deviant, the 145 ms deviant interestingly did not yield as substantial an effect as the 130 and 160 deviants. Because of the relatively crude determinations of the timeframes for the deviant conditions used in this experiment, further investigation that parses additional deviant variations to compare behavioral and neurological threshold levels would advance understanding of tactile temporal processing.

## Acknowledgements

The authors would like to acknowledge Kathryn Toffolo for sharing several analysis scripts that were helpful in the processing and visualization of this data, as well as Laura Ziemer, Emma Mantel, Eve Lang, Grace Rico, and Yacinda Hernandez for their assistance with data collection.

## Author Contributions

JX, ID-L, EF, and JF designed the study. JX and ID-L began data collection. EI collected the remaining data, processed and analyzed the data, and wrote the original draft of the manuscript. JX, ID-L, EF, and JF provided editorial input during manuscript preparation and editing. All authors read and approved the final version of the manuscript.

## Data Availability Statement

The authors will make the full de-identified dataset available (in EEG-BiDs format) through a public repository (e.g., DRYAD, Figshare) upon acceptance of this paper for publication, and will work with the editorial office to ensure that appropriate links are included in the final published version to allow for ease of access.

## Literature cited

1 Weber, A. I. et al. Spatial and temporal codes mediate the tactile perception of natural textures. Proc Natl Acad Sci U S A 110, 17107–17112 (2013). https://doi.org:10.1073/pnas.1305509110

2 Bensmaïa, S. J., Craig, J. C. & Johnson, K. O. Temporal factors in tactile spatial acuity: evidence for RA interference in fine spatial processing. J Neurophysiol 95, 1783–1791 (2006). https://doi.org:10.1152/jn.00878.2005

3 Gamzu, E. & Ahissar, E. Importance of temporal cues for tactile spatial-frequency discrimination. J Neurosci 21, 7416–7427 (2001). https://doi.org:10.1523/jneurosci.21-18-07416.2001

4 Hermosillo, R., Ritterband-Rosenbaum, A. & van Donkelaar, P. Predicting future sensorimotor states influences current temporal decision making. J Neurosci 31, 10019–10022 (2011). https://doi.org:10.1523/jneurosci.0037-11.2011

5 Yamamoto, S. & Kitazawa, S. Reversal of subjective temporal order due to arm crossing. Nature Neuroscience 4, 759–765 (2001). https://doi.org:10.1038/89559

6 Pöppel, E. A hierarchical model of temporal perception. Trends in Cognitive Sciences 1, 56–61 (1997). https://doi.org:https://doi.org/10.1016/S1364-6613(97)01008-5

7 Hirsh, I. J. & Sherrick, C. E., Jr. Perceived order in different sense modalities. J Exp Psychol 62, 423–432 (1961). https://doi.org:10.1037/h0045283

8 Mikkelsen, M. et al. Reproducibility of flutter-range vibrotactile detection and discrimination thresholds. Sci Rep 10, 6528 (2020). https://doi.org:10.1038/s41598-020-63208-z

9 Formby, C., Morgan, L. N., Forrest, T. G. & Raney, J. J. The role of frequency selectivity in measures of auditory and vibrotactile temporal resolution. J Acoust Soc Am 91, 293–305 (1992). https://doi.org:10.1121/1.402772

10 Näätänen, R., Paavilainen, P., Rinne, T. & Alho, K. The mismatch negativity (MMN) in basic research of central auditory processing: a review. Clin Neurophysiol 118, 2544–2590 (2007). https://doi.org:10.1016/j.clinph.2007.04.026

11 Näätänen, R., Gaillard, A. W. & Mäntysalo, S. Brain potential correlates of voluntary and involuntary attention. Prog Brain Res 54, 343–348 (1980). https://doi.org:10.1016/s0079-6123(08)61645-3

12 Ritter, W. & Ruchkin, D. S. A review of event-related potential components discovered in the context of studying P3. Ann N Y Acad Sci 658, 1–32 (1992). https://doi.org:10.1111/j.1749-6632.1992.tb22837.x

13 Novak, G., Ritter, W. & Vaughan, H. G., Jr. The chronometry of attention-modulated processing and automatic mismatch detection. Psychophysiology 29, 412–430 (1992). https://doi.org:10.1111/j.1469-8986.1992.tb01714.x

14 Ritter, W., De Sanctis, P., Molholm, S., Javitt, D. C. & Foxe, J. J. Preattentively grouped tones do not elicit MMN with respect to each other. Psychophysiology 43, 423–430 (2006). https://doi.org:10.1111/j.1469-8986.2006.00423.x

15 Alho, K., Woods, D. L., Algazi, A. & Näätänen, R. Intermodal selective attention. II. Effects of attentional load on processing of auditory and visual stimuli in central space. Electroencephalogr Clin Neurophysiol 82, 356–368 (1992). https://doi.org:10.1016/0013-4694(92)90005-3

16 Holopainen, I. E., Korpilahti, P., Juottonen, K., Lang, H. & Sillanpää, M. Abnormal frequency mismatch negativity in mentally retarded children and in children with developmental dysphasia. J Child Neurol 13, 178–183 (1998). https://doi.org:10.1177/088307389801300406

17 Ikeda, K., Hayashi, A., Hashimoto, S. & Kanno, A. Distinctive MMN relative to sound types in adults with intellectual disability. Neuroreport 15, 1053–1056 (2004). https://doi.org:10.1097/00001756-200404290-00024

18 Knoth, I. S. & Lippé, S. Event-related potential alterations in fragile X syndrome. Front Hum Neurosci 6, 264 (2012). https://doi.org:10.3389/fnhum.2012.00264

19 Nakajima, S. et al. Duration Mismatch Negativity Predicts Remission in First-Episode Schizophrenia Patients. Front Psychiatry 12, 777378 (2021). https://doi.org:10.3389/fpsyt.2021.777378

20 Haigh, S. M. et al. Mismatch negativity to pitch pattern deviants in schizophrenia. Eur J Neurosci 46, 2229–2239 (2017). https://doi.org:10.1111/ejn.13660

21 Javitt, D. C. Intracortical mechanisms of mismatch negativity dysfunction in schizophrenia. Audiol Neurootol 5, 207–215 (2000). https://doi.org:10.1159/000013882

22 Brima, T. et al. Auditory sensory memory span for duration is severely curtailed in females with Rett syndrome. Transl Psychiatry 9, 130 (2019). https://doi.org:10.1038/s41398-019-0463-0

23 Dunn, M. A., Gomes, H. & Gravel, J. Mismatch negativity in children with autism and typical development. J Autism Dev Disord 38, 52–71 (2008). https://doi.org:10.1007/s10803-007-0359-3

24 Ruiz-Martínez, F. J. et al. Impaired P1 Habituation and Mismatch Negativity in Children with Autism Spectrum Disorder. J Autism Dev Disord 50, 603–616 (2020). https://doi.org:10.1007/s10803-019-04299-0

25 Schwartz, S., Shinn-Cunningham, B. & Tager-Flusberg, H. Meta-analysis and systematic review of the literature characterizing auditory mismatch negativity in individuals with autism. Neurosci Biobehav Rev 87, 106–117 (2018). https://doi.org:10.1016/j.neubiorev.2018.01.008

26 Knight, E. J., Oakes, L., Hyman, S. L., Freedman, E. G. & Foxe, J. J. Individuals With Autism Have No Detectable Deficit in Neural Markers of Prediction Error When Presented With Auditory Rhythms of Varied Temporal Complexity. Autism Res 13, 2058–2072 (2020). https://doi.org:10.1002/aur.2362

27 Joutsiniemi, S.-L. et al. The mismatch negativity for duration decrement of auditory stimuli in healthy subjects. Electroencephalography and Clinical Neurophysiology/Evoked Potentials Section 108, 154–159 (1998).

28 Tales, A., Newton, P., Troscianko, T. & Butler, S. Mismatch negativity in the visual modality. Neuroreport 10, 3363–3367 (1999).

29 Czigler, I., Weisz, J. & Winkler, I. ERPs and deviance detection: visual mismatch negativity to repeated visual stimuli. Neuroscience letters 401, 178–182 (2006).

30 Kekoni, J. et al. Rate effect and mismatch responses in the somatosensory system: ERP-recordings in humans. Biological Psychology 46, 125–142 (1997). https://doi.org:https://doi.org/10.1016/S0301-0511(97)05249-6

31 Butler, J. S., Foxe, J. J., Fiebelkorn, I. C., Mercier, M. R. & Molholm, S. Multisensory representation of frequency across audition and touch: high density electrical mapping reveals early sensory-perceptual coupling. J Neurosci 32, 15338–15344 (2012). https://doi.org:10.1523/jneurosci.1796-12.2012

32 Spackman, L. A., Boyd, S. G. & Towell, A. Effects of stimulus frequency and duration on somatosensory discrimination responses. Exp Brain Res 177, 21–30 (2007). https://doi.org:10.1007/s00221-006-0650-0

33 Restuccia, D. et al. Somatosensory mismatch negativity in healthy children. Dev Med Child Neurol 51, 991–998 (2009). https://doi.org:10.1111/j.1469-8749.2009.03367.x

34 Shinozaki, N., Yabe, H., Sutoh, T., Hiruma, T. & Kaneko, S. Somatosensory automatic responses to deviant stimuli. Cognitive Brain Research 7, 165–171 (1998). https://doi.org:https://doi.org/10.1016/S0926-6410(98)00020-2

35 Shen, G., Weiss, S. M., Meltzoff, A. N. & Marshall, P. J. The somatosensory mismatch negativity as a window into body representations in infancy. Int J Psychophysiol 134, 144–150 (2018). https://doi.org:10.1016/j.ijpsycho.2018.10.013

36 Shen, G., Smyk, N. J., Meltzoff, A. N. & Marshall, P. J. Using somatosensory mismatch responses as a window into somatotopic processing of tactile stimulation. Psychophysiology 55, e13030 (2018). https://doi.org:10.1111/psyp.13030

37 Spackman, L. A., Towell, A. & Boyd, S. G. Somatosensory discrimination: An intracranial event-related potential study of children with refractory epilepsy. Brain Research 1310, 68–76 (2010). https://doi.org:https://doi.org/10.1016/j.brainres.2009.10.072

38 Chen, J. C. et al. Bi-directional modulation of somatosensory mismatch negativity with transcranial direct current stimulation: an event related potential study. J Physiol 592, 745–757 (2014). https://doi.org:10.1113/jphysiol.2013.260331

39 Akatsuka, K. et al. Mismatch responses related to temporal discrimination of somatosensory stimulation. Clinical Neurophysiology 116, 1930–1937 (2005). https://doi.org:https://doi.org/10.1016/j.clinph.2005.04.021

40 Strömmer, J. M., Tarkka, I. M. & Astikainen, P. Somatosensory mismatch response in young and elderly adults. Front Aging Neurosci 6, 293 (2014). https://doi.org:10.3389/fnagi.2014.00293

41 Butler, J. S. et al. Common or redundant neural circuits for duration processing across audition and touch. J Neurosci 31, 3400–3406 (2011). https://doi.org:10.1523/jneurosci.3296-10.2011

42 Mikkelsen, M., Wodka, E. L., Mostofsky, S. H. & Puts, N. A. J. Autism spectrum disorder in the scope of tactile processing. Dev Cogn Neurosci 29, 140–150 (2018). https://doi.org:10.1016/j.dcn.2016.12.005

43 De Sanctis, P., Molholm, S., Shpaner, M., Ritter, W. & Foxe, J. J. Right Hemispheric Contributions to Fine Auditory Temporal Discriminations: High-Density Electrical Mapping of the Duration Mismatch Negativity (MMN). Front Integr Neurosci 3, 5 (2009). https://doi.org:10.3389/neuro.07.005.2009

44 Delorme, A. & Makeig, S. EEGLAB: an open source toolbox for analysis of single-trial EEG dynamics including independent component analysis. J Neurosci Methods 134, 9–21 (2004). https://doi.org:10.1016/j.jneumeth.2003.10.009

45 Ide, M., Yaguchi, A., Sano, M., Fukatsu, R. & Wada, M. Higher Tactile Temporal Resolution as a Basis of Hypersensitivity in Individuals with Autism Spectrum Disorder. J Autism Dev Disord 49, 44–53 (2019). https://doi.org:10.1007/s10803-018-3677-8

46 Cascio, C. J. Somatosensory processing in neurodevelopmental disorders. J Neurodev Disord 2, 62–69 (2010). https://doi.org:10.1007/s11689-010-9046-3

47 Güçlü, B., Tanidir, C., Mukaddes, N. M. & Unal, F. Tactile sensitivity of normal and autistic children. Somatosens Mot Res 24, 21–33 (2007). https://doi.org:10.1080/08990220601179418

